# Systematic mapping of the free energy landscapes of a growing immunoglobulin domain identifies a kinetic intermediate associated with co-translational proline isomerization

**DOI:** 10.1101/189050

**Authors:** Christopher A. Waudby, Maria-Evangelia Karyadi, Tomasz Wlodarski, Anaïs M. E. Cassaignau, Sammy Chan, Julian M. Schmidt-Engler, Anne S. Wentink, Carlo Camilloni, Michele Vendruscolo, Lisa D. Cabrita, John Christodoulou

## Abstract

Co-translational folding is a fundamental molecular process that ensures efficient protein biosynthesis and minimizes the wasteful or hazardous formation of misfolded states. However, the complexity of this process makes it extremely challenging to obtain structural characterizations of co-translational folding pathways. Here we contrast observations in translationally-arrested nascent chains with those of a systematic C-terminal truncation strategy. We create a detailed description of chain length-dependent free energy landscapes associated with folding of the FLN5 filamin domain, in isolation and on the ribosome. By using this approach we identify and characterize two folding intermediates, including a partially folded intermediate associated with the isomerization of a conserved proline residue, which, together with measurements of folding kinetics, raises the prospect that neighboring unfolded domains might accumulate during biosynthesis. We develop a simple model to quantify the risk of misfolding in this situation, and show that catalysis of folding by peptidyl-prolyl isomerases is essential to eliminate this hazard.

## Introduction

The misfolding and aggregation of proteins is implicated in a wide range of human disorders [1]. It is therefore essential that cells have mechanisms to ensure quality control in protein synthesis and maintain protein homeostasis [2]. Increasing evidence indicates in particular that co-translational folding can increase the efficiency of folding when compared with the folding of full-length proteins *in vitro* [3]. Indeed, the ribosome is emerging as a hub in the cellular quality control network [4], regulating the interaction of nascent polypeptide chains with molecular chaperones and other factors, and in some cases directly modulating the folding process itself [5, 6].

The vast majority of protein folding studies to date have focused on small (≤150 residue) monomeric single-domain proteins, but such simple model systems are not fully representative of the bulk of the proteome [7]. A current challenge is therefore to apply our knowledge of fundamental folding mechanisms to more complex systems such as multi-domain proteins. Sequence analysis reveals that ca. 70% of eukaryotic proteins contain multiple domains, many of which are tandem repeat proteins, i.e. are composed of multiple copies of similar domains [8]. Such sequences are at high risk of forming non-native inter-domain contacts during the folding process, potentially leading to the formation of misfolded states [9, 10], and for this reason a strong selective pressure exists to minimize the homology between adjacent domains [11].

A major obstacle to the folding of multi-domain proteins is the exponential growth in the volume of conformational space that must be explored as the polypeptide chain length increases, which challenges the ability of a protein to fold readily and reliably. Co-translational folding is widely believed to play a major role in the solution of this problem, as the sequential folding of domains emerging from the ribosome exit tunnel restricts the volume of conformational space, thus promoting the rapid folding of the entire protein to its native state (while also minimising the risk of misfolding between adjacent domains). Therefore, in the context of the free energy landscape theory of protein folding, to study co-translational folding one should consider not a single free energy landscape, but a series of nested landscapes of increasing dimensionality and polypeptide chain lengths [12]. It is our goal here to map out these ‘co-translational landscapes’ in order to improve our understanding of the mechanisms by which the ribosome may contribute towards more efficient protein folding.

We have developed NMR spectroscopy as a tool for the observation of co-translational folding in translationally-arrested ribosome-nascent chain complexes (RNCs) [13–15], in particular to investigate the emergence and folding of a pair of immunoglobulin-like domains, FLN5-6, from the tandem repeat protein filamin (Fig. 1A,B) [6]. Folding of FLN5 to the native state was observed and a chemical shift-restrained ensemble structure was determined at a linker length well beyond the point at which the entire domain was solvent exposed (as probed by polyethylene glycol (PEG) mass-tagging assays).

**Figure 1.**
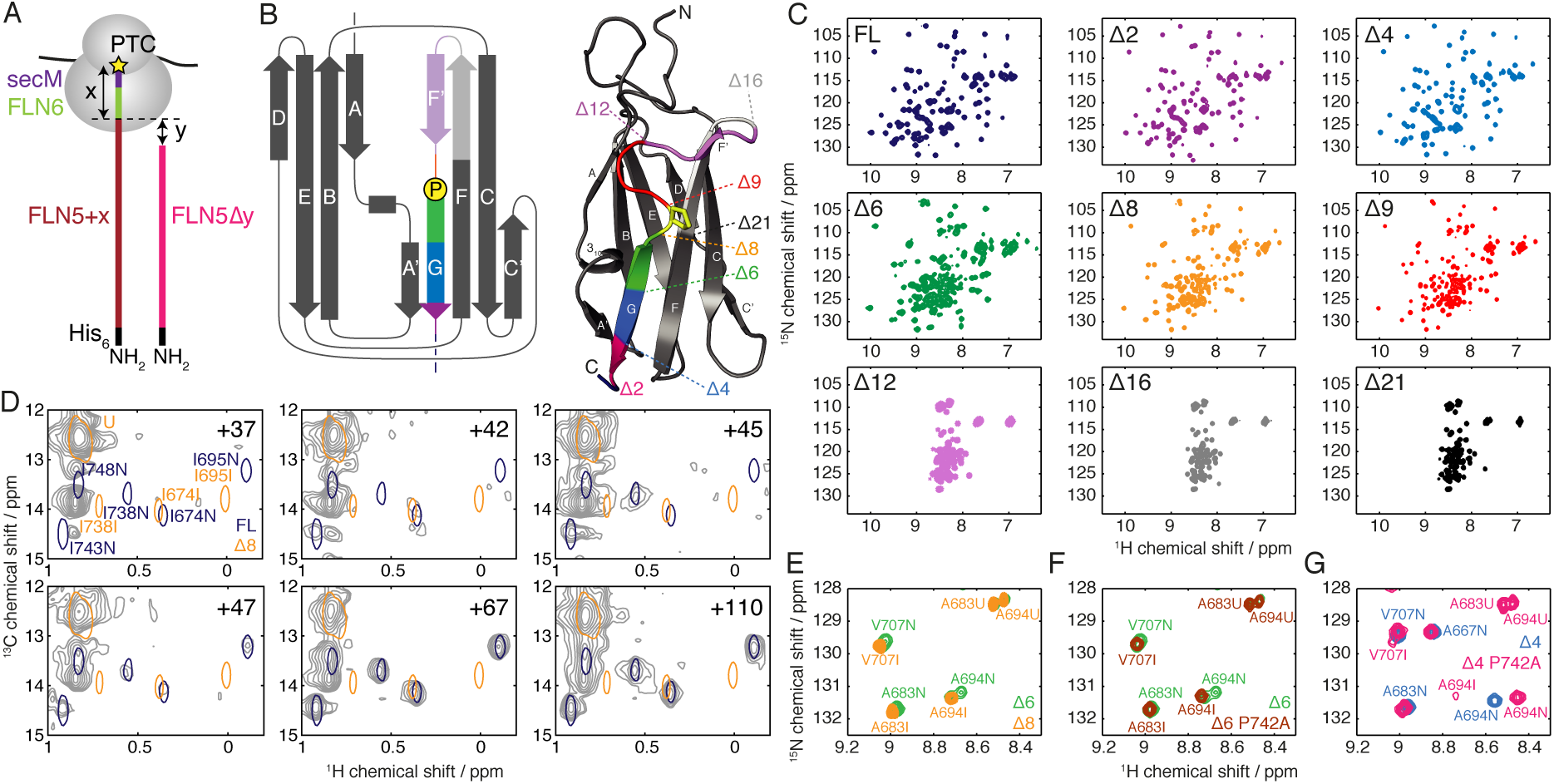
(A) Schematic and nomenclature of FLN5 RNC (FLN5+*x*) and truncation (FLN5Δ*y*) constructs. *x* and *y* represent the FLN6+secM linker length and the extent of truncation, respectively. (B) Topology and crystal structure of FLN5 (1qfh) colored and labeled to indicate the truncated constructs employed in this study. The *cis* proline P742 is highlighted in yellow (stick representation). (C) ^1^H,^15^N-SOFAST-HMQC spectra of FLN5 truncation variants (298 K, 600 MHz). (D) Comparison of ^1^H,^13^C-HMQC spectra of FL and Δ8 variants with spectra of [^13^CH_3_-Ile,^2^H]-labeled FLN5+*L* RNCs (gray) (298 K, 700 MHz). (E) Comparison of ^1^H,^15^N-HSQC spectra of FLN5Δ6 and FLN5Δ8, showing a magnified view of representative residues (283 K, 700 MHz). (F) Comparison of ^1^H,^15^N-HSQC spectra of FLN5Δ6 and FLN5Δ6(P742A) (283 K, 700 MHz). (G) Comparison of ^1^H,^15^N-HSQC spectra of FLN5Δ4 and FLN5Δ4(P742A) (283 K, 700 MHz).

Through a detailed analysis of a C-terminal truncation system, here we identify and determine the structure of an intermediate state associated with the isomerization of a conserved native-state *cis* proline. We then use this model to explore, and eliminate, the potential effects of steric occlusion in perturbing the folding process on the ribosome. By conducting a systematic exploration of the thermodynamics and kinetics of folding we create a detailed description of the free energy landscape for co-translational folding, from which the remarkably strong destabilizing effect of the ribosome may be quantified. Lastly, we present a simple model to analyze the effect of folding kinetics and proline isomerization on the risk of misfolding in this multi-domain protein.

## Results

Variants of the full-length (FL) FLN5 domain were designed to mimic the domain’s emergence during translation by truncating between 2 and 21 residues from the C-terminus (denoted Δ2 to Δ21), corresponding to deletions from the G strand up to the F strand (Fig. 1A,B). Biochemical characterization using native polyacrylamide gel electrophoresis (Fig. S1A) showed Δ12 and Δ16 to have expanded conformations relative to FL, while the migration of Δ6 to Δ9 was approximately midway between these states. A transition from *β*-sheet to random-coil structure was also observed at these lengths by far-UV circular dichroism (CD) spectroscopy (Fig. S1B).

To investigate this length-dependent unfolding transition in greater detail, 2D NMR correlation spectra of truncation variants were acquired (Figs. 1C and S1C). The large proton chemical shift dispersion observed in FL, Δ2 and Δ4 spectra indicates the presence of stable tertiary structure, while the narrow chemical shift dispersion observed in spectra of the Δ12, Δ16 and Δ21 variants is characteristic of a disordered state. At intermediate lengths (Δ6, Δ8 and Δ9) multiple sets of cross-peaks were observed, indicating that both structured and unstructured states are populated in equilibrium, interconverting in slow exchange on the NMR timescale (approximately seconds or slower). Two states, designated ‘I’ (intermediate) and ‘U’ (unfolded), were populated by Δ8 and Δ9 variants, while an additional set of folded resonances (‘N’, native-like) was observed in the Δ6 spectrum (Fig. 1E, S2).

We hypothesized that the two sets of folded resonances in the Δ6 spectrum arise from *cis*/*trans* isomerization of the native-state *cis* proline P742 (Fig. 1B). To test this conjecture we examined the Δ6 P742A variant, in which P742 was mutated to shift the equilibrium towards the non-native *trans* conformation. The additional resonances observed in the Δ6 spectrum were absent in spectra of Δ6 P742A (Fig. 1F), while the remaining well-dispersed resonances corresponded with those observed in the I state of the Δ8 variant, in which P742 is the C-terminal residue, and the Δ9 variant in which P742 is absent altogether. We therefore conclude that there is a length-dependent transition from an intermediate state ‘I’ in which P742, if present, is in a *trans* conformation, to a (near-)native state ‘N’ in which P742 is in a *cis* conformation. Consistent with this conclusion, in the Δ4 P742A variant (Fig. 1G) the N state was destabilized and both I and U resonances could additionally be observed with the same chemical shifts as observed in WT Δ6.

The structured states of the FLN5 nascent chain can be observed with the greatest sensitivity using a specific isoleucine C*δ* methyl labeling strategy, the chemical shifts of which are very sensitive to the local chemical environment and hence folded structure [6]. However, no resonances corresponding to the intermediate state were detected in any of these RNC spectra (Fig. 1D). We can therefore conclude that the intermediate observed in isolation is either structurally incompatible with tethering to the ribosome, or is destabilized by some other mechanism. To distinguish between these possibilities we therefore sought to understand the intermediate structure in more detail.

### Structural characterization of folding pathway

We investigated the structure of C-terminal truncation variants by first monitoring progressive changes in amide chemical shifts from the FL native state to the Δ6 I state, which were found to be clustered around the A and F strands adjacent to the truncated G strand (Fig. S3). The magnitude of chemical shift perturbations increased with the extent of truncation up to the Δ6 I state, but no large chemical shift differences were observed in shorter lengths indicating that the intermediate structure does not change significantly beyond this point.

We then used chemical shift-restrained replica-averaged metadynamics simulations [16] to determine ensemble structures of the FL native and Δ6 intermediate states (Fig. 2A-B and S4). The quality of the ensembles was comparable to structures previously determined by chemical shift methods [17], assessed by back-calculated chemical shift deviations, residual dipolar coupling Q factors (Table S1), and a backbone RMSD of 1.8 Å between the FL ensemble and a crystal structure [18] (Fig. 2A). We find that the FL ensemble is generally well ordered, with C*α* RMSF values below 0.5 Å except in some loop regions (Fig. 2A).

**Figure 2.**
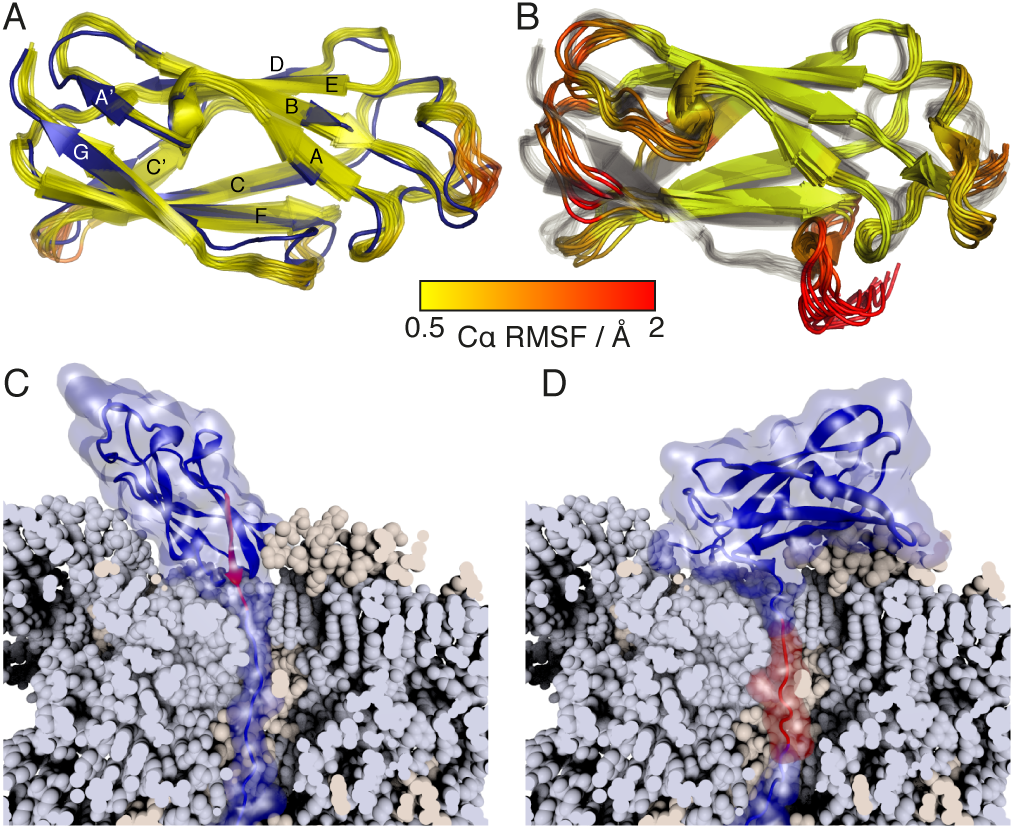
Structural analysis of FLN5 truncation variants. (A) Ensemble structure of FL FLN5 (yellow–red), aligned against the previously determined crystal structure (1qfh, blue) [18]. The FL ensemble is colored according to the C*α* RMSF as indicated by the key below. (B) Ensemble structure of the Δ6 *trans* intermediate (colored by the C*α* RMSF), aligned against the FL ensemble (gray). (C,D) Modeling of the closest possible approach of (C) native and (D) intermediate FLN5 structures tethered to the ribosome (shown in a cut-away view to highlight the NC path through the exit tunnel). The G strand of FLN5 (disordered in the intermediate) is highlighted in red.

In contrast, in the Δ6 intermediate state (Fig. 2B) a native-like core is retained, comprising the A–F strands, but the truncated G strand is unfolded, with increased disorder also found in the adjacent A’ and F strands. No structure could be determined for the Δ6 N state, due to extensive broadening of resonances around the site of the truncation (Fig. S5A). ^15^N CPMG relaxation dispersion measurements [19] of adjacent residues indicate this arises from conformational exchange on a millisecond timescale (Fig. S5B). We suggest that this exchange may reflect the transient association of the truncated G strand with the F strand, which is then stabilized as the polypeptide chain length increases.

In these relaxation measurements, small dispersions (*R*_ex_ < 4 s^−1^) were also observed in the unfolded state of Δ6 at 283 K (Fig. S6A). These dispersions could be fitted as a two-state exchange process, with ^15^N chemical shift differences of up to 8 ppm between this high-energy intermediate state, populated to 0.95 *±* 0.05 % and designated ‘I*’, and the unfolded ground state (Fig. S6B). Chemical shift differences were uniformly small (< 0.5 ppm) for residues 646–670, but from residue 671 onwards they were correlated with those between Δ6 U and I states, indicating the formation of intermediate-like structure in this region. Thus, the I* intermediate comprises a native-like core formed from the B to F strands, while both the A strand and the truncated G strand remain disordered (Fig. S6C). The intermediate was also identified in the Δ9, Δ12 and Δ16 variants, with a decreased population, 0.4 *±* 0.1 %, in the shortest length, Δ16 (Fig. S6D–I).

In summary therefore, the length-dependent folding pathway proceeds from the unfolded state, via a high energy intermediate with disordered A and G strands, to a stable intermediate with a disordered G strand and P742, if present, in a *trans* conformation. Isomerization of P742 to *cis* then allows association of the G strand to form the native state.

Considered in the context of co-translational folding, the disordered C-terminal G strand in the intermediate suggested that this state should be structurally compatible with formation as an emerging ribosome-bound nascent chain. To explore this further we ran a series of simulations using structure-based models [20] with various linker lengths to study folding of the N and I states on the ribosome. Using this approach, we found that the native state requires a 20 aa linker (in an extended conformation) to fold on the ribosome, whereas the linker length needed for folding to the intermediate is only 14 aa (Fig. 2C,D). By comparison with the linker conformation in an all-atom MD simulation of the FLN5+110 RNC [6], we can further estimate that these native and intermediate structures are accessible with linker lengths (in a relaxed conformation) of ca. 26 and 18 aa respectively (Fig. S7). The crucial observation from this modelling is therefore that there is no steric reason that the intermediate should not be populated earlier than the native state during biosynthesis, and that formation of the intermediate must be specifically inhibited by ribosomal tethering.

### Thermodynamic analysis

We next sought to complement our structural description of the folding pathway during elongation with a detailed energetic description of the co-translational folding landscape. The stabilities of FL, Δ2 and Δ4 variants were measured by chemical denaturation, while in Δ6, Δ8 and Δ9, where multiple states were observed to be populated at equilibrium, NMR measurements of ^1^H,^15^N-HSQC peak volumes were used to determine the relative populations of folded and unfolded states and hence calculate free energies of folding directly (Fig. 3 and S8). Where only the unfolded state could be observed (Δ12 and Δ16), limits on the stability of the intermediate have been calculated based on observed signal-to-noise ratios (dashed lines). From these measurements, we found that the six C-terminal residues contribute almost the entire stability of the domain, Δ*G*_N−D_ = −7.0 ± 0.2 kcal mol^−1^. The stability of the I state did not vary significantly between Δ6 and Δ9 (Δ*G*_I−U_ = 0.8 ± 0.2 kcal mol^−1^), but in shorter lengths the intermediate was further destabilized, with a population below the ca. 1% limit of detection (derived from signal-to-noise levels).

**Figure 3.**
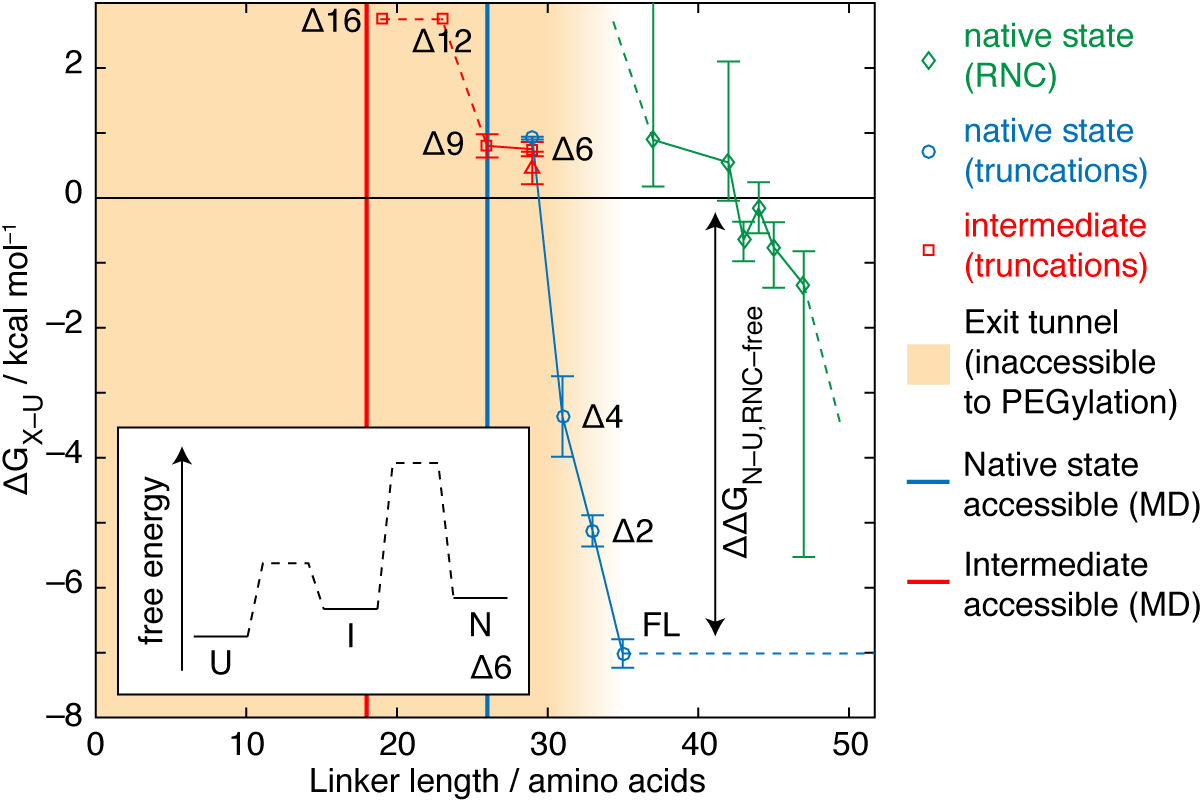
Thermodynamic characterization of the FLN5 co-translational folding pathway. Free energies of folding to native (*cis*-P742) and intermediate (*trans*-P742) states at 298 K are shown, in truncated variants and RNCs as indicated. As a specific example the Δ6 folding pathway is illustrated in the inset figure. A red triangle indicates Δ6 intermediate stability as determined by magnetization transfer measurements (Fig. 4A,B). Yellow shading indicates the extent of the ribosome exit tunnel, as determined by PEG accessibility [6], which was used as the basis for alignment of the isolated length-dependent and ribosome-associated folding pathways, while solid lines indicate the points at which native and intermediate states were found to be accessible in MD simulations (Fig. 2C,D).

In a similar manner to the truncated variants, we then used NMR peak volumes to determine the chain length dependence of folding in RNCs. As the linewidths of folded methyl resonances varied strongly with chain length, affecting the observed intensities or volumes, only unfolded state resonances that showed uniform linewidths across all chain lengths were used in the analysis (Fig. 3). The alignment of truncation and RNC chain lengths is based upon the complete accessibility to PEG observed for residues 36 aa from beyond the PTC [6], and therefore we assigned the FL isolated state to this length. We note that this is a conservative alignment when compared with our structure-based modeling (Fig. 2C,D), which predicts that linker lenghts of ca. 18 and 26 aa are required for folding to the intermediate and native states respectively (Fig. 3, solid lines). In either case, it is clear that the ‘delay’ in NC folding following emergence from the exit tunnel must be associated with a strong destablization of the NC, with a ΔΔ*G* comparable to the 7 kcal mol^−1^ folding energy of the isolated domain.

### Measurement of folding kinetics

Co-translational folding is a non-equilibrium process, with folding occurring in parallel with elongation of the NC. It is therefore important to understand the rate of NC folding in addition to the thermodynamic characterization above. Folding and proline isomerization typically occur over a wide range of timescales, which require a range of spectroscopic techniques to resolve. 2D N_*z*_-exchange measurements [21] of Δ6 (Fig. 4A) showed exchange cross-peaks between unfolded and intermediate states, but no exchange with the native state, indicating that the timescale of exchange was much longer than the nitrogen *T*_1_ (ca. 1 s). Where I and N resonances could be clearly resolved (e.g. for A694 but not A683, Fig. 4A), peak intensities were measured as a function of the mixing time and these were fitted globally to determine the rates of folding and unfolding (Fig. 4B). For Δ6 at 298 K, we found the folding rate *k*_UI_ = 0.8 ± 0.2 s^−1^, and the unfolding rate *k*_IU_ = 1.7 ± 0.4 s^−1^. This corresponds to a free energy for folding Δ*G*_I−U_ = 0.4 ± 0.3 kcal mol^−1^, consistent with our earlier measurement of 0.8 *±* 0.2 kcal mol^−1^ based on integration of peak volumes (Fig. 3, red triangle).

**Figure 4.**
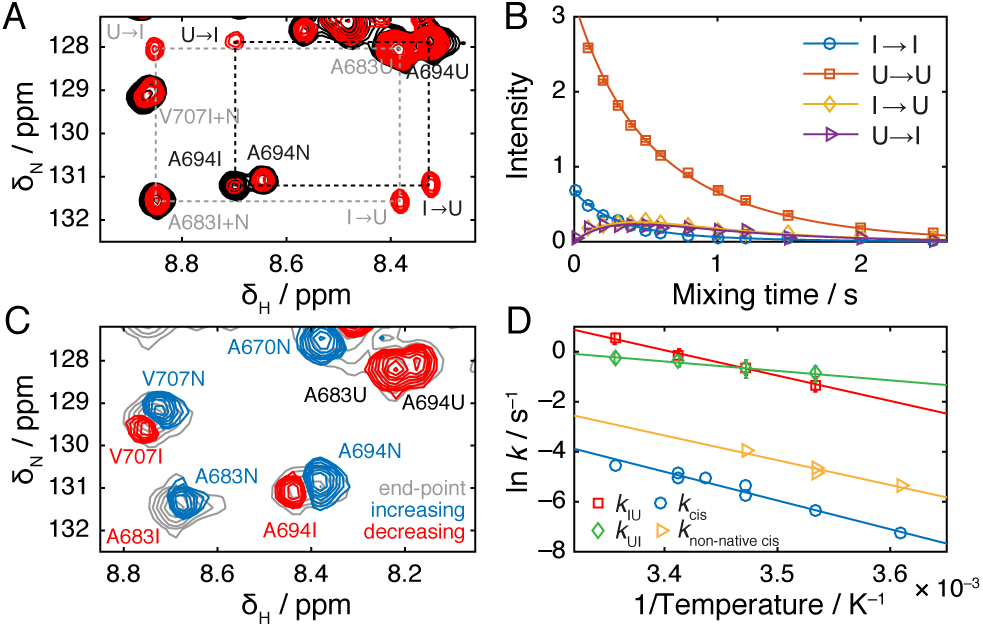
Analysis of folding kinetics. (A) 2D slices from a N_*z*_ magnetization transfer experiment [21] at 298 K, with exchange times of 10 ms (black) and 500 ms (red). Dashed rectangles highlight exchange cross-peaks between unfolded and intermediate states. Grey coloring indicates peaks excluded from further analysis due to overlap of native and intermediate resonances. (B) Analysis and fitting of A694 cross-peak intensities from the magnetization transfer experiment shown in (A). Data from multiple resonances were fitted globally to determine the indicated exchange rates. (C) Component spectrum determined from the analysis of real-time NMR data for the refolding of Δ6 following a temperature jump from 310 K to 298 K, with a fitted time constant of 1.5 min. Peaks that increase or decrease over the reaction time-course are shown in blue and red respectively. The spectrum at equilibrium is shown in gray. (D) Arrhenius plot showing the temperature dependence of Δ6 folding kinetics, with fitted activation energies as indicated. *k*_IU_ and *k*_UI_ are exchange rates determined by magnetization transfer measurements; *k*_non−native−cis_ and *k*_cis_ are proline isomerization rates determined by real-time NMR methods.

To measure the proline-isomerization limited rate of Δ6 folding to the native state, we developed a real-time NMR measurement of relaxation kinetics following a rapid temperature jump from a thermally unfolded state at 310 K (Figure S1) to refolding conditions between 277 and 298 K. A series of ^1^H,^15^N-SOFAST-HMQC spectra [22] were acquired rapidly (15 s per spectrum) until equilibrium was re-established, and these spectra were fitted globally to a constant plus one or two exponential phases. The resulting amplitude spectra contain a combination of positive and negative peaks, describing resonances that increase and decrease over time respectively (Figure S9A).

For the refolding of Δ6 at 298 K, *trans*-to-*cis* isomerization of P742 was observed with a time constant of 1.5 min (Figure 4C). At lower temperatures an additional faster phase was also identified that we attribute to off-pathway *cis*-to-*trans* isomerization of seven native-state *trans* prolines in the unfolded state (Figure S9B). Consistent with this interpretation, *trans*-to-*cis* isomerization was not observed in measurements of Δ6 P742A refolding (Figure S9C), while the refolding of Δ4 P742A at 283 K fitted to a single slow phase with a time constant of 17.3 min, indicating formation of a *cis* alanine (Figure S9D). Finally, activation energies of 21.7 *±* 1.0 and 20.0 *±* 0.7 kcal mol^−1^ were determined for the fast and slow phases of Δ6 refolding (Figure 4D), comparable with typical activation energies for proline isomerization (20–22 kcal mol^−1^) [23].

### Reconstructing the non-equilibrium co-translational folding pathway

In this work, we have analyzed the thermodynamics and kinetics of FLN5 folding as a function of polypeptide chain length, and have identified states that are structurally compatible with tethering to the ribosome during translation (Figures 2–4). This description can be combined with a known translation rate (ca. 5 amino acids s^−1^ in eukaryotes) to construct a Markov chain model of the co-translational folding pathway (Figure S10), which then can be integrated numerically to obtain quantitative predictions about the fundamentally non-equilibrium co-translational folding process [24]. From this model, we observe significant differences between populations calculated at thermodynamic equilibrium and those predicted under non-equilibrium conditions (Figure 5A). In particular, as translation is more rapid than *trans*-*cis* isomerization (i.e. folding to the native state), only 17% of FLN5 NCs are predicted (based on our analysis of isolated truncations) to be fully folded by the point at which the subsequent FLN6 domain has been translated. Such an accumulation of unfolded domains may present a misfolding hazard, explored further in the next section.

**Figure 5.**
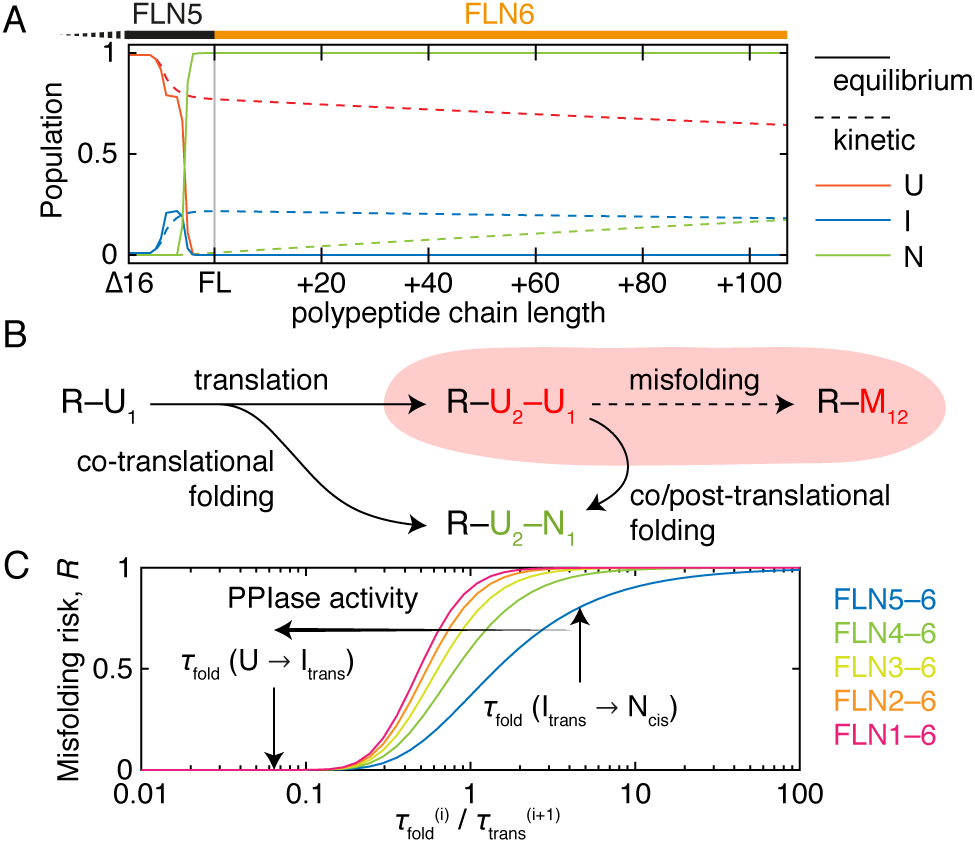
Reconstruction of the non-equilibrium co-translational folding pathway and analysis of the risk of misfolding during protein synthesis. (A) Populations of folded, unfolded and intermediate states, based on characterization of isolated truncations, are shown as a function of polypeptide chain length at equilibrium (i.e. infinitely slow/stalled translation, solid lines) and under non-equilibrium conditions with a translation rate of 5 amino acids s^−1^ (dashed lines). (B) A model for assessing the risk of co-translational misfolding, in which the rate of folding of a ribosome-bound unfolded NC (R–U_1_) to its native state (N_1_), relative to the rate at which the following domain (U_2_) is translated, determines the likelihood of populating adjacent unfolded domains (R–U_1_–U_2_) which may be at risk of forming a misfolded state (M_12_). (C) Estimated risk of misfolding in filamin, as a function of the number of tandem repeat domains and the relative rates of folding and translation. Arrows indicate predicted risks where folding is limited by proline isomerization to the native state (right), or, through the action of peptidyl-prolyl isomerases, limited by folding to the intermediate (left).

### Assessing the risk of co-translational misfolding in tandem repeat proteins

Filamins are tandem repeat proteins comprising six immunoglobulin domains (in *D. discoideum*, and 24 in the case of human filamins A, B and C), and the sequence identity between adjacent domains is 31 *±* 7 % (mean *±* s.d.) with 9% of pairs above the 40% threshold associated with misfolding in other immunoglobulin domains (Table S2) [11]. Moreover, the proline at the start of the G strand is present and conserved in a *cis* conformation in every available structure (50% coverage of *D. discoideum* and human filamin domains, Table S2). Thus, the co-translational accumulation of adjacent unfolded domains predicted above results not only in an explosive increase in the volume of accessible conformational space, that co-translational folding phenomena were postulated to help avoid, but introduces a risk of misfolding interactions between adjacent domains [9, 10, 25].

To investigate this further we developed a simple model to quantify the risk of misfolding (Figure 5B). By analysing the timescale of co-translational folding, 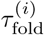, relative to translation of the following domain, 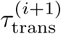, we find (Supporting Information) that for a protein with *N* tandem repeat domains, the misfolding risk, *R*, given by the probability of populating adjacent unfolded domains during biosynthesis, is 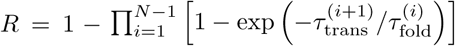.

This expression is plotted in Figure 5C for the FLN5-6 pair studied here directly, and also for the entire tandem repeat region, given a translation rate of 5 aa s^−1^ (although we expect our results to be robust with respect to small variations in this rate). In isolation, when folding is limited by proline isomerization (Fig. 5C, right-hand arrow), the misfolding risk is high, particularly when the full multi-domain protein is considered. If, however, this step is accelerated enzymatically, by peptidyl-prolyl isomerases, the folding time will tend towards a new limit given by the rate of folding from the unfolded state (Fig. 5C, left-hand arrow). Under this condition, we find that the risk of misfolding is essentially eliminated.

## Discussion

During protein synthesis on the ribosome, free energy landscapes for NC folding evolve as the chain length is increased by translation [12]. Characterization of a co-translational folding process therefore requires the characterization of a series of free energy landscapes, rather than the single surface associated with traditional folding studies. Here we have begun this process, using observations of translationally-arrested RNCs, under equilibrium conditions, complemented by a systematic C-terminal truncation strategy [26–28], to explore these landscapes with structural (Figure 2), thermodynamic (Figure 3) and kinetic (Figure 4) detail.

The truncation model we have developed in this work has two important roles. Firstly, it provides an opportunity to apply high resolution spectroscopic methods that are not currently feasible directly on the ribosome, to analyze structure and folding kinetics. In doing so we have identified a folding intermediate associated with the isomerization of a native-state *cis* proline. Secondly, the model provides a baseline against which the effects of the ribosome itself may be discerned, highlighting the large free energy changes (ΔΔ*G* ∼ 7 kcal mol^−1^) associated with the ‘delay’ of co-translational folding by only a small number of residues (Fig. 3). We have previously attributed this effect to stabilization of the unfolded state by interactions with the ribosome surface, driven by high effective concentrations when the nascent chain is tethered near the exit tunnel [6]; a key challenge for future work will be to develop a quantitative understanding of this holdase activity.

Due to the dynamic nature of the truncated intermediates and the extensive resonance overlap and exchange between states, truncation intermediates were not amenable to structural analysis using conventional crystallographic or NOE-based methods. Therefore, we employed chemical shift-restrained replica-averaged metadynamics simulations [16] to calculate an ensemble structure of the intermediate (Figure 2B). This structure, having a disordered G strand, was used as a starting configuration for structural modelling of the intermediate as a nascent chain, from which we concluded that there are no steric barriers to its formation co-translationally. Instead, we suggest that the same interactions of the unfolded state with the ribosome surface that inhibit formation of the native state also act at shorter chain lengths to destabilize the intermediate.

It is interesting to compare the intermediate we identify here with the aggregation-prone intermediate formed during folding, or upon N-terminal truncation, of *β*_2_-microglobulin [29,30]. Both domains have an immunoglobulin fold, and both intermediates are associated with the isomerization of a native-state *cis* proline (P32 in the case of *β*_2_-microglobulin). However, a key difference is that FLN5 is part of a multi-domain protein, and so the primary hazard is the intramolecular misfolding of adjacent unfolded or intermediate states during translation, rather than post-translational intermolecular aggregation. Such misfolded states have been found to form ubiquitously between neighbouring domains, and where sequence identity is high their formation may become effectively irreversible [9, 10, 25]. We have developed a simple model to assess this risk (Figure 5B,C), and found that PPIase activity is essential to eliminate this risk (although it is interesting to consider why such an apparent folding hazard should be so strongly conserved at all (Table S2)). Critically, our findings highlight that thermodynamically favourable co-translational folding is not by itself sufficient to avoid misfolding hazards: kinetic requirements must also be satisfied.

## Materials and Methods

### Protein expression and purification

FLN5 truncations were prepared by site-directed mutagenesis (Stratagene) by replacing the residue of choice with a stop codon. Isotopically unlabeled and uniformly ^15^N, ^13^C/^15^N and ^2^H/^13^C/^15^N labeled proteins were overexpressed in *E. coli* BL21-GOLD cells (Stratagene) and purified as previously described [6]. Purified proteins were exchanged into Tico buffer (10 mM HEPES, 30 mM NH_4_Cl, 6 mM MgCl_2_, 0.02% sodium azide, 1 mM EDTA, pH 7.5). 1 mM BME was also included in samples of full-length FLN5 and the Δ2 truncation to inhibit formation of disulphide-bonded dimers at Cys747.

### Sequence analysis

Multiple sequence alignments were generated using the MATLAB Bioinformatics Toolbox (2016b, The MathWorks Inc., MA) using the BLOSUM80 scoring matrix. Percent identities are reported relative to the number of non-gap positions.

### CD spectroscopy

CD spectra of truncation variants were acquired at 283 K using a Jasco J-810 spectropolarimeter (Jasco Corporation, Tokyo), with protein concentrations of ca. 0.5 mg mL^−1^ and urea concentrations varied between 0 and 8.2 M. Samples were equilibrated for 1–4 hours in a water bath at 283 K and CD signals at 234 nm were then measured as a function of the denaturant concentration and globally fitted to a two-state unfolding model using a shared *m*-value, in order to determine the free energy of folding.

### NMR spectroscopy

Unless otherwise noted, NMR data were acquired at 283 K on a 700 MHz Bruker Avance III spectrometer equipped with a TXI cryoprobe, using uniformly ^15^N or ^15^N/^13^C labeled protein at concentrations of 0.2–1 mM in Tico buffer supplemented with 10% D2O (v/v) and 0.001% DSS (w/v) as an internal chemical shift reference. Data were processed with nmrPipe and cross-peak intensities were measured using FuDA (http://www.biochem.ucl.ac.uk/hansen/fuda/).

### Resonance assignment

FL, Δ4, Δ6 and Δ9 FLN5 spectra were assigned by standard triple-resonance methods. ^1^H,^15^N difference spectra obtained from real-time NMR measurements of Δ6 refolding were used to assist the assignment of closely overlapping resonances arising from *cis* and *trans* intermediates. Additionally, a 0.2 mM sample of uniformly ^2^H/^15^N-labelled Δ6 was used to record a 2D N_*z*_ magnetization exchange spectrum [31] (mixing time 1 s) to cross-validate the assignment of unfolded and intermediate resonances. Only small chemical shift changes were observed between the unfolded and intermediate states of Δ6 and Δ9, and cross-peak assignments of Δ9 were therefore transferred directly from those of Δ6 and cross-validated using a 2D N_*z*_ magnetization transfer measurement. The assignments of ^1^H,^15^N-HSQC resonances of unfolded Δ12 and Δ16 variants were similarly transferred from the complete Δ6 and Δ9 assignments.

### Intensity analysis

HSQC spectra were acquired of uniformly 15N-labeled proteins at concentrations of 300–350 *µ*M and processed with exponential apodization in both dimensions. Resonances were fitted with Lorentzian lineshapes and observed volumes were corrected for the effects of transverse relaxation during INEPT transfer periods [32]. For each spectrum, Δ*G*_a−b_ = −*RT* ln(*V*_a_*/V*_b_) was calculated independently for several residues using cross-peaks that were well resolved in all states, and stabilities have been reported as the mean *±* s.e.m. of these measurements.

In Δ12 and Δ16 ^1^H,^15^N-HSQC spectra, the only resonances that could be detected were those arising from the U state, and therefore it is only possible to place a limit on the maximum stability using the signal-to-noise ratio in the spectrum to estimate the maximum possible population of unobserved structured state. The observed signal-to-noise ratios in Δ12 and Δ16 spectra were between 250 and 300, but to reflect the additional uncertainty arising from possible changes in the position and line-widths of folded cross-peaks, the limit of detection was capped at 1%. From this, a minimum Δ*G*_*I*−*U*_ of 2.7 kcal mol^−1^ was calculated.

RNC stabilities were calculated as a function of linker length using the intensity of selected ^1^H,^15^N-SOFASTHMQC resonances, measured relative to the partially emerged and fully unfolded +21 RNC [6]. As the selected resonances were previously shown not to exhibit length-dependent changes in linewidth [6], their relative intensity was used directly as a measure of the unfolded state population and used to determine free energies of folding. Uncertainties were propagated using a Monte Carlo approach.

### Magnetization transfer experiments

2D N_*z*_ longitudinal magnetization transfer experiments [31] were acquired for Δ6, Δ6 P742A and Δ9 with magnetization transfer delays between 0.01 and 2.5 s. The intensities of residues for which complete sets of exchange cross-peaks could be observed between folded (F) and unfolded (U) states were fitted to a numerical solution of the exchange process:

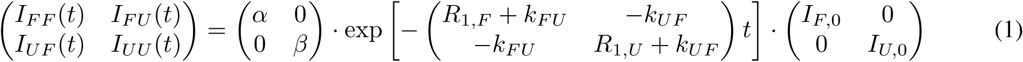

where *I_FU_* is the amplitude of the F-to-U cross-peak etc., *I*_*F*,0_ and *I*_*U*,0_ are the amplitudes of the folded and unfolded states at the start of the magnetization transfer period, and *α* and *β* are factors accounting for relaxation following this period. Data for multiple residuals were fitted globally with shared exchange rates *k*_*FU*_ and *k*_*UF*_. Uncertainties in the fitted exchange parameters were determined by bootstrap resampling of residuals.

### Real-time temperature-jump kinetics

Real-time NMR temperature-jump measurements were acquired for samples of Δ6, Δ6 P742A and Δ4 P742A. Cooling gas was provided using a BCU-05 unit with a flow rate of 935 L hr^−1^. The time required for the temperature within NMR samples to re-equilibrate after lowering the temperature from 310 K was measured by repeated acquisition of 1D ^1^H NMR spectra of a sample of d_4_-methanol during a programmed temperature change, and we found that the internal temperature stabilized to within 1 K within 80–120 s depending on the final temperature (277–298 K). We observed little difference in the rate of temperature change in 3 mm vs 5 mm diameter NMR tubes, and therefore 5 mm shigemi tubes were used to maximize sensitivity while minimizing formation of temperature gradients within the sample.

^1^H,^15^N-SOFAST-HMQC experiments [22] (2 scans, 1024 × 32 complex points, 15.7 s per spectrum) were prepared and calibrated with a sample incubated at the target temperature. The sample temperature was then raised to 310 K and equilibrated for at least 15 min to induce unfolding. We note that the extrapolated timescale of cis/trans isomerization at this temperature is 20 s (Figure 4D), so the system is expected to be fully equilibrated under these conditions. The repeated acquisition of 2D experiments was then begun and a temperature change was initiated. Acquisition was continued for 1–2 hr until the re-equilibration process was complete. This process was repeated a number of times to increase the signal to noise ratio of the final data. Sample integrity during these temperature cycles was monitored using 1D ^1^H spectra and the first increments of the 2D spectra.

Experimental time-courses were fitted globally to a constant plus one or two exponential phases using a variable projection algorithm [33]. After Fourier transformation in the direct dimension and extraction of the amide region, the pseudo-3D data *Y*_*mij*_(*τ*, *t*_1_, *ω*_2_) were reshaped into an *M* × *N* matrix *y*_*mn*_, where *M* is the number of time points and *N* the total number of points in each 2D spectrum. We then attempted to find solutions of the form *y* = *CA* + *ϵ*, where *C* is an *M* × *L* matrix describing the changing concentration of spectral components over time, *A* is a *L* × *N* matrix describing the spectral amplitudes of these components, and *ε* is the error term. *C* is parameterized by the rate constants *k*_*l*_ for each kinetic process, *C*_*ml*_(*k*) = exp(−*k*_*l*_τ_*m*_), and we fix *k*_1_ = 0 to represent the equilibrium end-point. Thus, we attempt to find min_*A*,*k*_ ‖*C*(*k*)*A* − *y*‖_*F*_. This is a difficult non-linear optimization problem in a high-dimensional space, but if *k* is fixed then the solution is simply *A* = *C*^+^(*k*)*y*, where *C*^+^ is the Moore-Penrose pseudoinverse. By exploiting this conditionally linear structure, the initial problem can be reduced to min_*k*_ ‖*C*(*k*)*C*^+^(*k*)*y* − *y*‖_*F*_. This is the far simpler problem of non-linear minimization in the rates alone, which we solve using a Trust-Region-Reflective algorithm. Reported uncertainties in the rate constants are calculated from the Jacobian approximation to the covariance matrix. The amplitudes *A* of the component spectra can then be found by projection, and these are reshaped back into 2D spectra and Fourier transformed in the indirect dimension for further visualization and analysis.

### CPMG relaxation dispersion

^15^N CW-CPMG relaxation dispersion experiments [19] were acquired at 500 and 700 MHz. Uniformly ^15^N-labelled samples of Δ9, Δ12 and Δ16 were prepared, while measurements of Δ6 were conducted using uniform ^2^H/^15^N-labelling. Experiments were performed at 283 K, using a ca. 10 kHz ^1^H spin-lock, 5.56 kHz ^15^N CPMG refocusing pulses, and a 40 ms relaxation delay with ^1^H and ^15^N temperature compensation elements. Changes in the sample temperature during acquisition were measured using the changes in the H_2_O chemical shift and the external temperature was adjusted to maintain the desired internal temperature. Dispersion profiles were fitted to two-state exchange models using CATIA http://www.biochem.ucl.ac.uk/hansen/catia/) in a two step process: firstly, residues with large exchange contributions were fitted globally to determine the global exchange parameters (*k*_ex_ and *p_B_*); these parameters were then held constant and used to fit chemical shift differences for residues showing weaker dispersions.

### Structure determination

Structures of the ground states were obtained from chemical shift-restrained replica-averaged metadynamics [16] performed using the Amber99SB*-ILDN force field. All the simulations were run in GROMACS 4.6.5 using PLUMED 1.3 and Almost to introduce metadynamics and chemical shift restraints. The particle-mesh Ewald method was used for long-range electrostatic interactions, with a short range cut-off of 0.9 nm, and for Lennard-Jones interactions a 1.2 nm cut-off was used. Simulations were run in the canonical ensemble using velocity rescaling with a stochastic term for temperature coupling [34]. Simulations were carried out for FL and Δ6 I (with P742 in a trans conformation). Initial structures for each of the systems were prepared from the crystal structure, 1qfh [18], with addition of N-terminal hexahistidine tags, truncation of the C-terminus as required, and changing the isomeric state of P742 to *trans* in the Δ6 model. After initial equilibration runs, each system was simulated for ca. 450 ns in four replicas in parallel at 283 K, with a harmonic restraint applied on the average value of CamShift back-calculated chemical shifts [35]. In addition, each replica was sampling one of the four collective variables (CVs) applied to: function of the *φ* and *ψ* dihedral angles (ALPHABETA1), the *χ*_1_ and *χ*_2_ dihedral angles for hydrophobic and polar amino acids (ALPHABETA2), the fraction of the antiparallel *β*-sheet conformations (ANTIBETARMSD), and the radius of gyration (RGYR). Exchanges between replicas were attempted periodically every 50 ps according to a replica-exchange scheme and well-tempered scheme [36], with a bias-factor of 8.0 used to rescale the added bias potential. Convergence of the simulations was reached after 350 ns which resulted in free energy landscapes with statistical uncertainties of ≤2.0 kJ mol^−1^ for free energies up to 10 kJ mol^−1^ (Figure S3g-h). Analysis of the metadynamics trajectory and the assignment of microstates was carried out in VMD using the METAGUI plugin. Structural representatives of each ground state were chosen from the lowest free energy microstate using criteria of the lowest MolProbity score. Residual dipolar couplings were back-calculated from structural ensembles using PALES and ensemble averaged values were used to calculate quality factors, 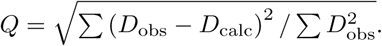. Hexahistidine purification tags were excluded from analyses of RMSD/RMSF and are not shown in figures.

Folding simulations on the ribosome for I and N states were conducted in Gromacs 4.6.5 with all-atom structure-based models generated in SMOG 2 [20]. Simulated systems combined a fragment of the ribosome around the exit tunnel and the FLN5 domain in an extended conformation tethered to the PTC via linker with variable length from 15 aa to 31 aa. The linker sequence was the same as used in FLN5 RNCs studies [6]. Ribosome atoms were frozen during the simulations, whereas for FLN5 contacts from N or I states were applied to simulate folding; linker residues were not restrained.

### Markov chain modeling

Transition matrices were calculated by linear interpolation of measured stabilities as a function of chain length. Rate constants were interpolated assuming that folding rates are approximately independent of chain length, with the unfolding rate being modulated according to the measured stability, *k*_*u*_ = *k*_*f*_ exp(Δ*G*/*RT*). We also assumed that folding to the native state occurs directly from the intermediate, although given the separation in the timescales of folding and proline isomerization, I and U are effectively in pre-equilibrium and the impact of this choice is minimal. Populations (folding probabilities) were then calculated as a function of time using the matrix exponential of the transition matrix, given an initial population of Δ16 in the unfolded state. Populations were also calculated as a function of polypeptide chain length by insertion of absorbing states after each translation step to quench further reactions, an approach equivalent to that of O’Brien et al. [24].

### Calculation of misfolding risk

Following synthesis of domain *i*, the expected translation time of the following domain is 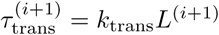, where *k*_trans_ is the translation rate and *L*^(*i*+1)^ is the number of residues in domain *i* + 1. As the sum of a series of identical exponential processes (i.e. averaging out sequence-specific variations in *k*_trans_), *τ*_trans_ has an Erlang distribution with rate *k*_trans_ and shape parameter *L*. *τ*_trans_ therefore has a narrow distribution, with coefficient of variation *L*^−1/2^, and so will be approximated as a constant hereonin. The probability of domain *i* folding during translation of domain *i* + 1 is then 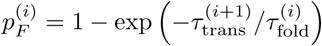, where 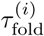 is the time constant for folding, in the present case given by 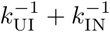 (approximating folding with single exponential kinetics). Therefore, for a protein with *N* tandem repeat domains, the misfolding risk, *R*, given by the probability of populating adjacent unfolding domains during biosynthesis, is 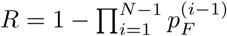.

## Supporting Information

**Table S1.** NMR and refinement statistics for protein structures.

**Table S2.** Sequence identity (with preceding domain) and structural analysis of conserved proline conformation across filamin domains.

**Figure S1.** Biochemical and spectroscopic characterization of truncation variants.

**Figure S2.** ^1^H,^15^N HSQC spectrum (283 K, 700 MHz) of FLN5Δ6 showing resonance assignments.

**Figure S3.** Amide chemical shift changes, relative to FL FLN5.

**Figure S4**. Free energy landscapes determined by chemical shift-restrained replica-averaged metadynamics.

**Figure S5.** Millisecond timescale dynamics in the FLN5 Δ6 *cis* state.

**Figure S6.** CPMG relaxation dispersion measurements of unfolded states.

**Figure S7.** Comparison of NC linker conformations in structure-based models of the native and intermediate states, with an all-atom, chemical shift-restrained molecular dynamics simulation of the FLN5+110 RNC.

**Figure S8.** Thermodynamic characterization of folding in FLN5 truncation variants at 283 K.

**Figure S9.** Real-time NMR measurement of proline isomerization kinetics following temperature jumps.

**Figure S10.** Markov chain modelling of the non-equilibrium co-translational folding pathway.

## Acknowledgments

We are grateful to Dr Flemming Hansen for helpful discussions, and to Dr John Kirkpatrick for assistance with NMR experiments. We acknowledge the use of the UCL NMR Centre and Legion High Performance Computing Facility (Legion@UCL), the MRC for access to the Biomedical NMR Centre at the Francis Crick Institute, London, and the staff for their support. Funding was provided by the Wellcome Trust, TW was supported by an EMBO fellowship, and MK and SC by BBSRC studentships.

